# Experimental Snowball Earth Viscosity Drives the Evolution of Motile Multicellularity

**DOI:** 10.1101/2024.02.06.579218

**Authors:** Andrea Halling, Brysyn Goodson, Anna Hirschmann, Boswell A. Wing, Carl Simpson

## Abstract

During the 70-million-year span of the Cryogenian Snowball Earth glaciations, low ocean temperatures beneath global sea ice increased water viscosity up to fourfold. In the absence of adaptation, unicellular organisms living in this viscous environment were limited in their ability to move and acquire nutrients. We experimentally test the hypothesis that multicellularity evolved in order to overcome this viscosity-induced metabolic deficit. In the presence of Snowball Earth viscosities, we find that populations of unicellular green algae evolve motile multicellular phenotypes in addition to other phenotypes that optimize different combinations of size and speed. As the Snowball Earth subsided and warm seas returned, the novelty of motile multicellularity permitted these organisms to take physical control over their local environment for the first time. This innovation may underpin the evolution of dominant multicellular lineages on Earth today.

**Significance statement:** Beginning 720-million years ago, two global glaciations — together known as the Snowball Earth — covered the planet with a thick layer of ice for a total of 70-million years. Several groups of complex multicellular organisms independently radiated at this time, including animals, green algae, and red algae. All of these clades include lineages with large bodies made of thousands of cells, multiple cell types, and spatial organization. At first glance, it seems that life merely survived despite the Snowball Earth glaciations. We find experimental evidence that the Snowball Earth glaciations were instead an evolutionary trigger for the diversification of complex multicellular groups.

## Introduction

Simple eukaryotic multicellularity has evolved many times in Earth’s history (1–3) and easily evolves in response to numerous experimental selective pressures in the lab (4–8). While this experimental and paleontological work has helped demystify the process, it highlights a paradox in our understanding: if multicellularity is so easy to evolve, why did modern multicellular lineages that have dominated ecosystems for the last 540 million years take so long to diversify after the origin of eukaryotes? Investigating the temporally unique geological selective pressures when dominant multicellular lineages radiated can help to resolve this paradox.

All sources of information on the origins of obligately multicellular eukaryotes are biased in different ways, but the direction of those biases paints a clear picture (9). Molecular clock dates — which tend to be biased toward older dates (10–13) — indicate that the origins of animals occurred within the Tonian, prior to the Snowball Earth Glaciations (14, 15). Early fossilized animals – which tend to be biased toward more recent dates (11–13) – appear in the Cryogenian interglacial interval (16) as well as the Ediacaran after the Snowball Earth glaciations are over (Fig. 1, 16, 17). The molecular clock dates of multicellular green and red algae are clustered in the Tonian-Cryogenian (18, 19). Although some fossils of unclear modern affinity appear earlier in the rock record, small filamentous multicellular algae become common in the Tonian (3, 20–24). In the Cryogenian interglacial Nantou flora, larger algal fossils appear with a significant increase in surface area to volume ratio (20, 25). Collectively, these sources indicate that the early history and radiation of multicellular clades coincide temporally with the Tonian-Cryogenian Snowball Earth interval (Fig. 1).

**Figure 1:**
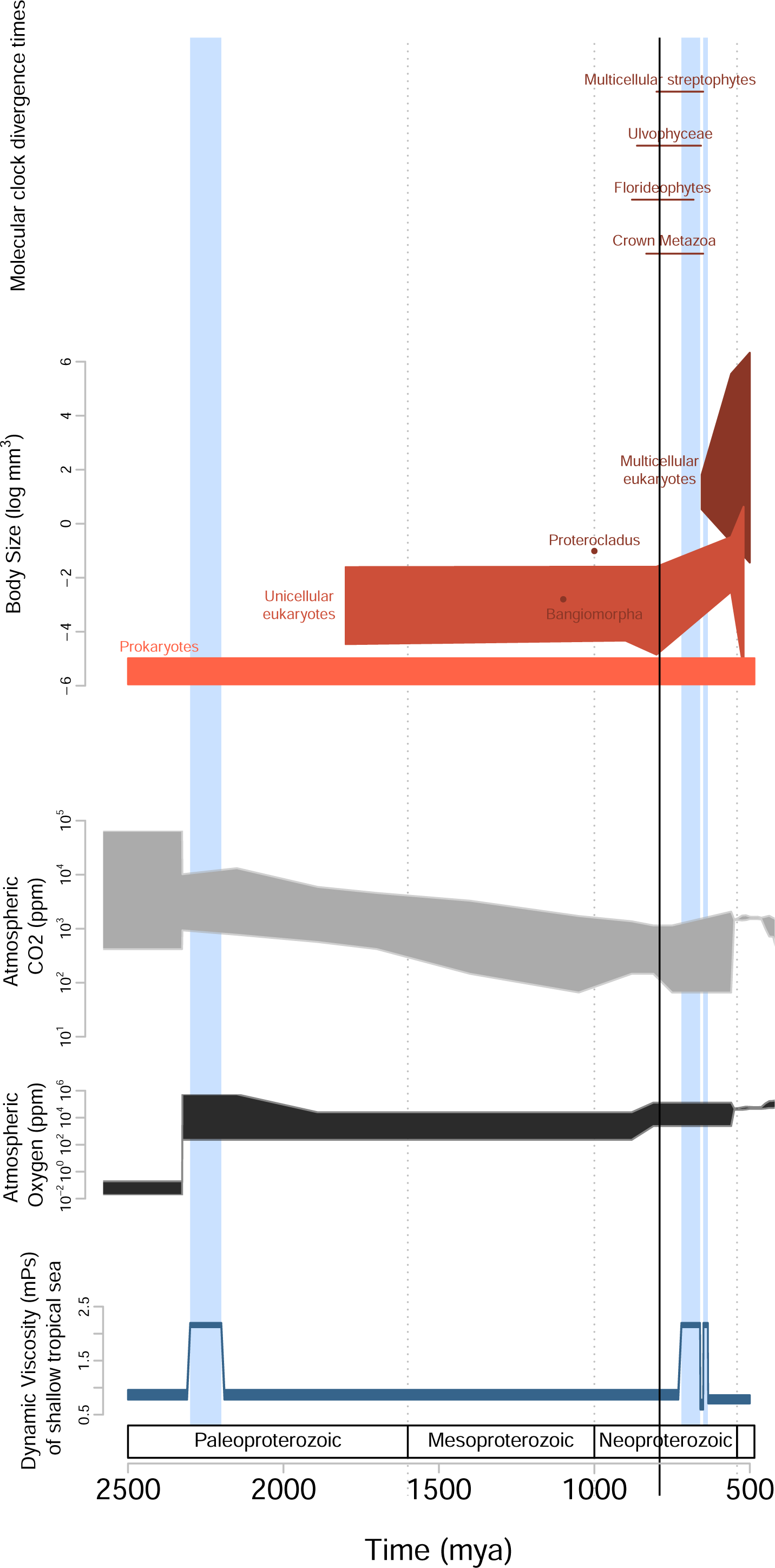
Schematic evolution of the Earth System over the billion years of Proterozoic time. 95% confidence intervals for molecular clock divergence time estimates for multicellular lineages (18, 19, 70). Body volume for fossils are split out by prokaryotes (bacteria and archaea), unicellular eukaryotes, and multicellular eukaryotes (17, 55). The earliest multicellular eukaryotic algae, *Protocladus* (21) and *Bangiomorpha* (71) are called out separately. Proxies and modeling provide estimates for CO2 ppm (72–74)(60, 61, 63) and O2 ppm (73–75). Inferred seawater viscosity for shallow equatorial ocean with glacial interval viscosities for seawater with temperatures from −4 to −3 °C and salinities of 35 ppt. Interglacial seawater is shown with 35 ppt seawater and temperatures ranging from 25 - 35 °C (55).

The oxygen requirement of modern multicellular organisms has long lead to the inference that oxygen levels increased along with the rise of animals and other multicellular organisms (2, 26, 27). We now know that the significant rise of atmospheric oxygen levels occurred after the Cambrian explosion (28–33) nearly 200-million years too late for it to be the cause of the origin of multicellularity in animals and algae. In addition to this mismatch in timing, high seawater temperatures in the aftermath of the Snowball Earth glaciations would have physiologically limited access to oxygen (34–36). Therefore, the ability for animals to use any increasing oxygen levels would have been suppressed by thermal stress until the oceans cooled sufficiently (37). Furthermore, it has been shown that high levels of oxygen actually suppress the size of experimentally evolved multicellular organisms (38) complicating any mechanism that directly links oxygen to large size.

It is also hypothesized that continental weathering caused by the Snowball Earth glaciations (39, 40) resulted in a massive increase in nutrient availability selecting for multicellularity, despite a likely much earlier occurrence of high nutrient levels (41). Cryogenian nutrient influx does well to explain the shift from bacterial primary productivity to productivity dominated by eukaryotes (39, 42), but does not easily explain the origin of multicellularity, even if it helps explain the later Cambrian explosion (43). Biotic drivers, such as predation and grazing, are also a common hypothesis for the selection of multicellular eukaryotes (5–7, 44). Yet eukaryotic predation evolved with heterotrophy during eukaryogenesis (45–47) and predates the origin of multicellularity (47, 48). Moreover, unicellular autotrophs often possess the ability to curtail predator-prey arms races (49).

Prior to the Cryogenian, life would have likely lived predominately in shallow, warm tropical seas (50). As early as 780 mya (51, 52), seawater temperature drastically decreased as the oceans became uniformly cold (53). For organisms adapted to living in warm seawater, this would have become a homogeneous global selective filter. This temperature decrease would result in a global increase of water viscosity (Fig. 1). Life could survive under thick sea ice but to access food, organisms would have had to contend with the high viscosity of the seawater.

With little observed extinction (54) and no spatial refugia (53), populations of Cryogenian unicellular eukaryotes must have adapted to the cold viscous conditions of Snowball Earth in order to overcome the metabolic deficit imposed on them. There are several hypothesized strategies that unicellular organisms could have used in order to minimize this deficit by optimizing size, motility, and metabolic scaling (55, 56). Unicells could have become smaller, lowering their metabolic rates to match availability.

Alternatively, unicells could have increased nutrient acquisition by becoming multicellular. Larger sizes together with higher collectively generated speeds would have increased encounter rates enough to satisfy metabolic needs of larger bodies (55). Evolution driven by the physical changes of seawater during the Snowball Earth has the potential to explain both the timing and the exclusively eukaryotic nature of complex multicellularity (55, 56).

To test the biological feasibility of this hypothesis, we isolate the physical aspect of viscosity to use as a selective regime in microbial evolution experiments. Our experimental petri dish (macroplate, Fig. S1), inspired by Baym et al 2016 (57), utilizes a stepwise viscosity gradient. This macroplate spatially represents the temporal increase in viscosity as Snowball Earth conditions set in. Our setup provides a stable gradient over the timescale of weeks to months as diffusion of our viscosity-inducing polymer (Ficoll400) occurs slowly, as a function of length squared. We inoculate wildtype *Chlamydomonas reinhardtii* — a unicellular green alga — at the standard viscosity (1x) of HSA media in the center of the circular Macroplate. As the population grows out radially, only individuals with the ability to move into the sections of higher viscosity (2− and 4x relative viscosities) continue to propagate. The spatially structured environment of the Macroplate acts as a vehicle for experimental evolution, with viscosity as the selective barrier, and nutrient availability as the evolutionary driver.

## Results

We observe that *C. reinhardtii* evolves a diversity of phenotypic responses to media of 2x and 4x relative viscosities. This includes flagellated motile colonies of four and eight cells, small non-motile unicells, and macroscopic non-motile colonies. Coulter Counter size distributions as well as microscopy of the populations living within the 2x and 4x sections of the macroplate reflect this phenotypic shift (Fig. 2).

**Figure 2:**
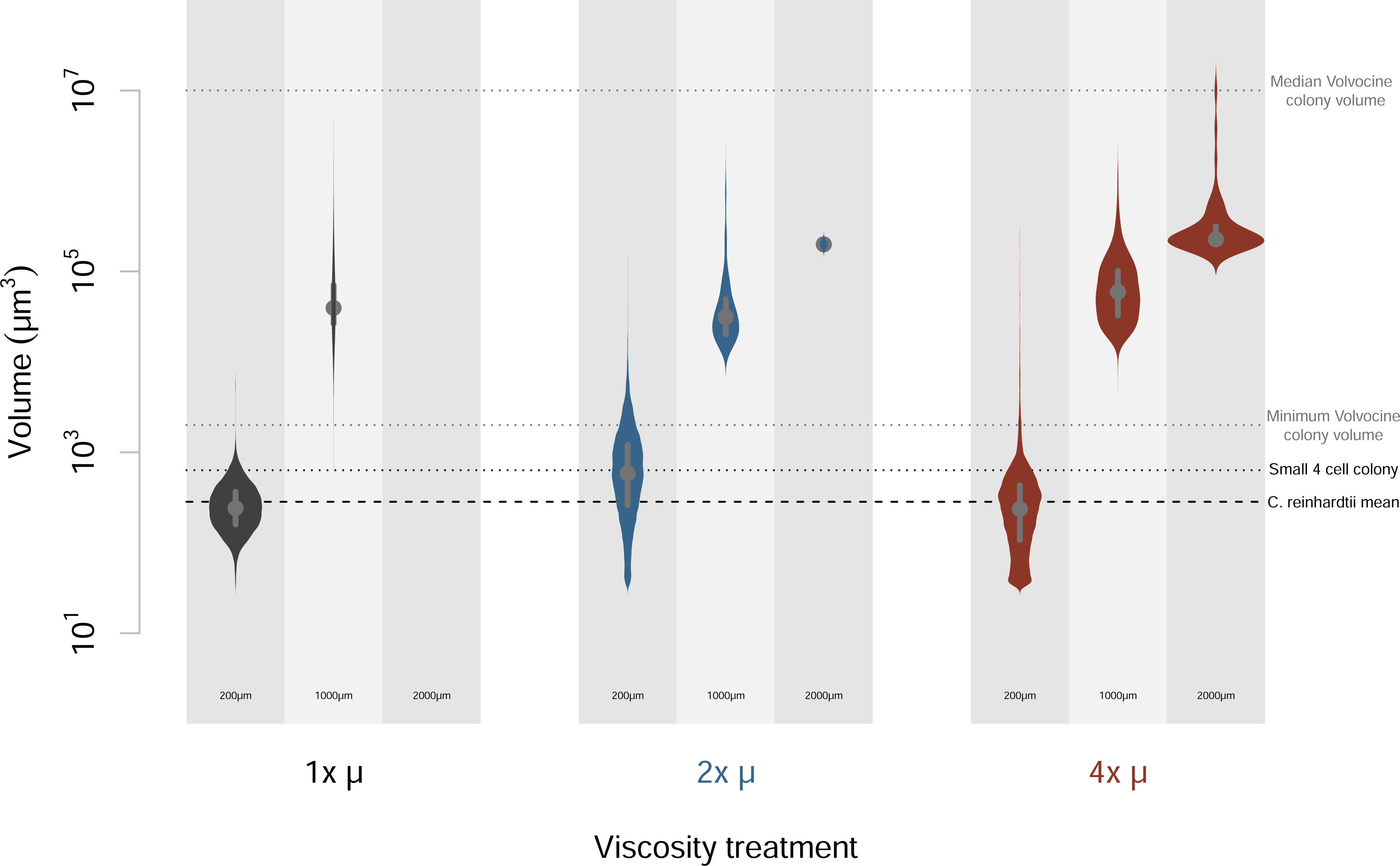
Size distributions of plastic and evolved populations. Sizes of >13,000 individuals from each plastic population as well as from each radial section from the macroplate. The left side of each density plot (light gray) shows the non-evolved, phenotypic plastic response of *C. reinhardtii* to different levels of viscosity. The right side of each density plot shows the evolved response of *C. reinhardtii* in each viscosity from the macroplate. Separate size distributions are plotted for three sizes of coulter counter aperture tube: 200µm, 1000µm, and 2000µm. For reference, the range and median volume for colonial volvocales (76, 77) are shown as horizontal reference lines.

In addition to unique size distributions, seemingly coordinated motility was observed in four and eight-celled individuals from both 2x and 4x relative viscosities (Fig. 3). As expected, (55, 58), single cells decreased in speed as viscosity increased (∼12 µm s^-1^ in 1x, ∼4 µm s^-1^ in 2x, and ∼1.7 µm s^-1^ in 4x relative viscosities) while colonies of four and eight cells maintain average speeds near ∼15µm s^-1^ in 2x and 4x relative viscosities (Movie S1). These motile phenotypes are maintained even when returned to media of 1x viscosity and transferred for at least 70 generations (Fig. 4).

**Figure 3:**
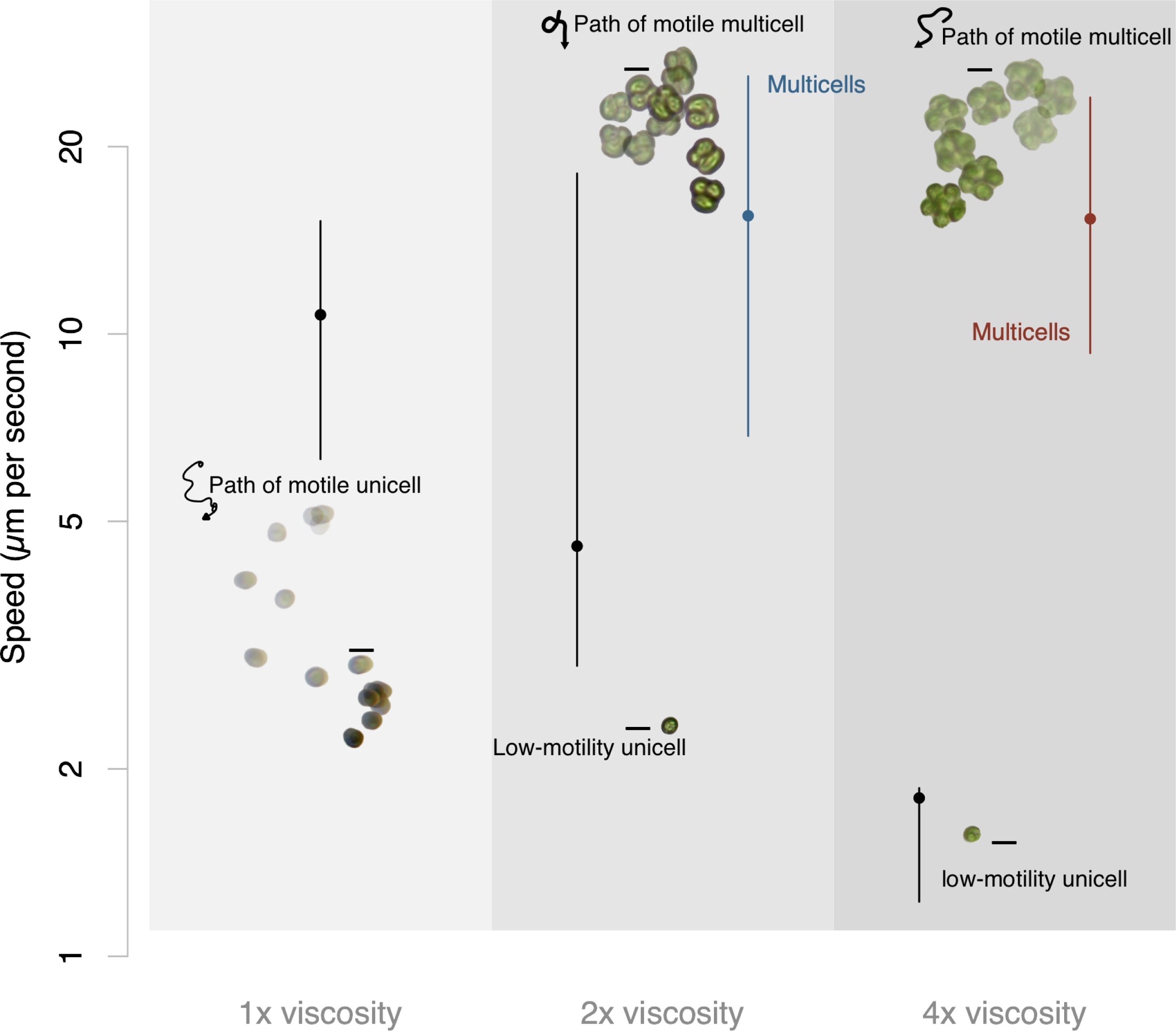
Quantification of motility in evolved populations. Microscopy videos analyzed by custom cell tracking code were used to find speeds of evolved individuals. Black data points show the decreasing average speeds of single cells from the macroplate in their corresponding increasing viscosities. Blue shows the average speed of motile multicells in 2x relative viscosity and red shows the average speed of motile multicells in 4x relative viscosity Examples of motile phenotypes from evolved populations in 1x, 2x, and 4x relative viscosities. Motile unicellular phenotypes were the dominant morphology in the 1x viscosity section of the macroplate. When viscosity increased to 2x relative viscosity, motile multicells of four emerged. In the 4x relative viscosity section of the macroplate, motile multicells of both four and eight cells were abundant in the population. Motility is shown as a stop-frame animation with about 10 frames between each image and all phenotypes are displayed on the same scale.

**Figure 4:**
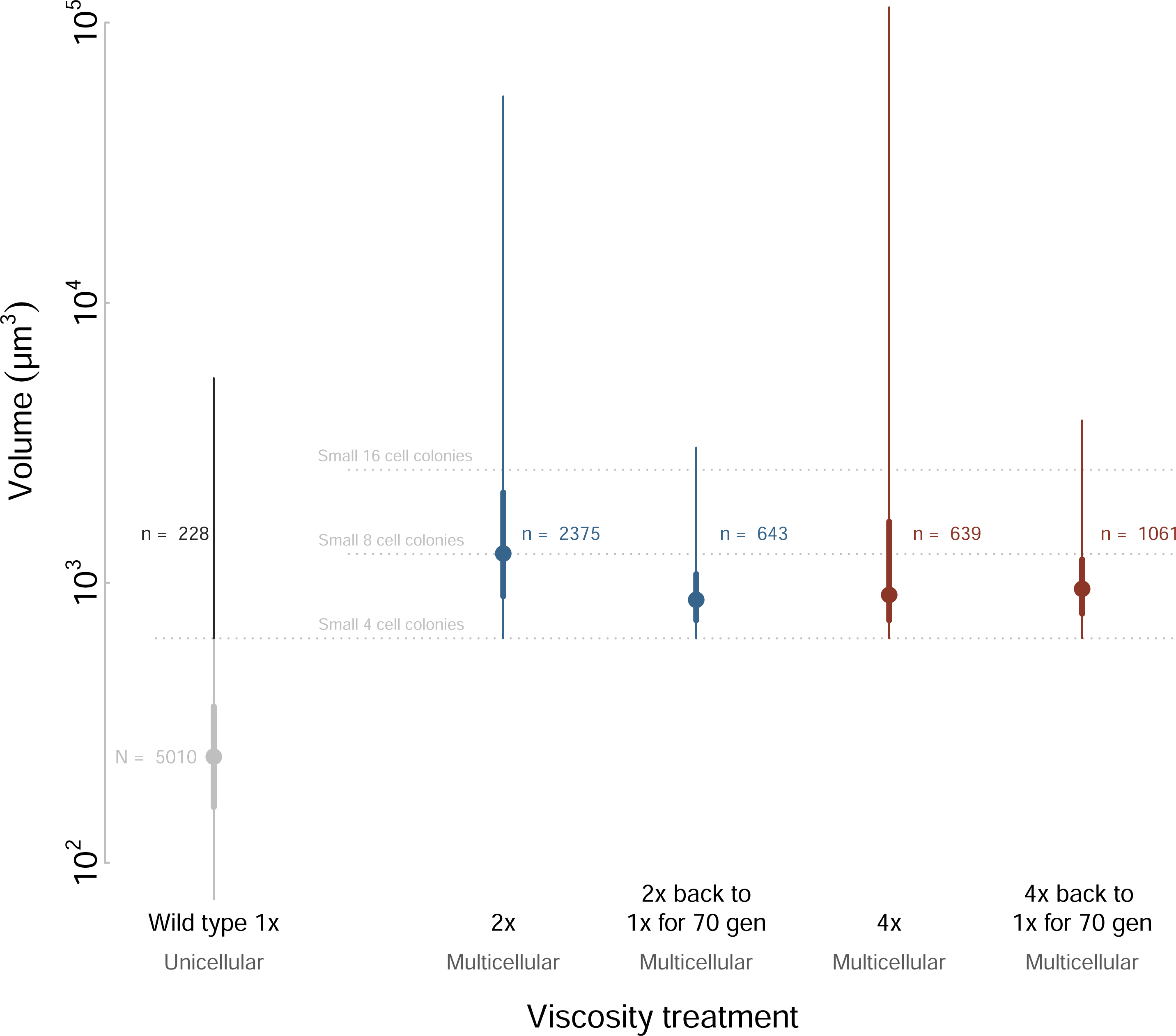
Size distributions of heritability in evolved populations. Wildtype *C. reinhardtii* in 1x viscosity is compared with evolved populations when returned to 1x media and maintained for over 75 generations. The left side of each density plot (light gray) represents the wild-type size distribution. These distributions reflect the return of plastic populations to typical wild-type distributions when placed back into 1x media. The right side of each density plot shows the size distributions of evolved 2x and 4x relative viscosity populations when returned to 1x relative viscosity media. Gray bars show the size range of multicells containing four and eight cells. Motile multicellular phenotypes continued to be abundant in low viscosity.

**Figure 5:**
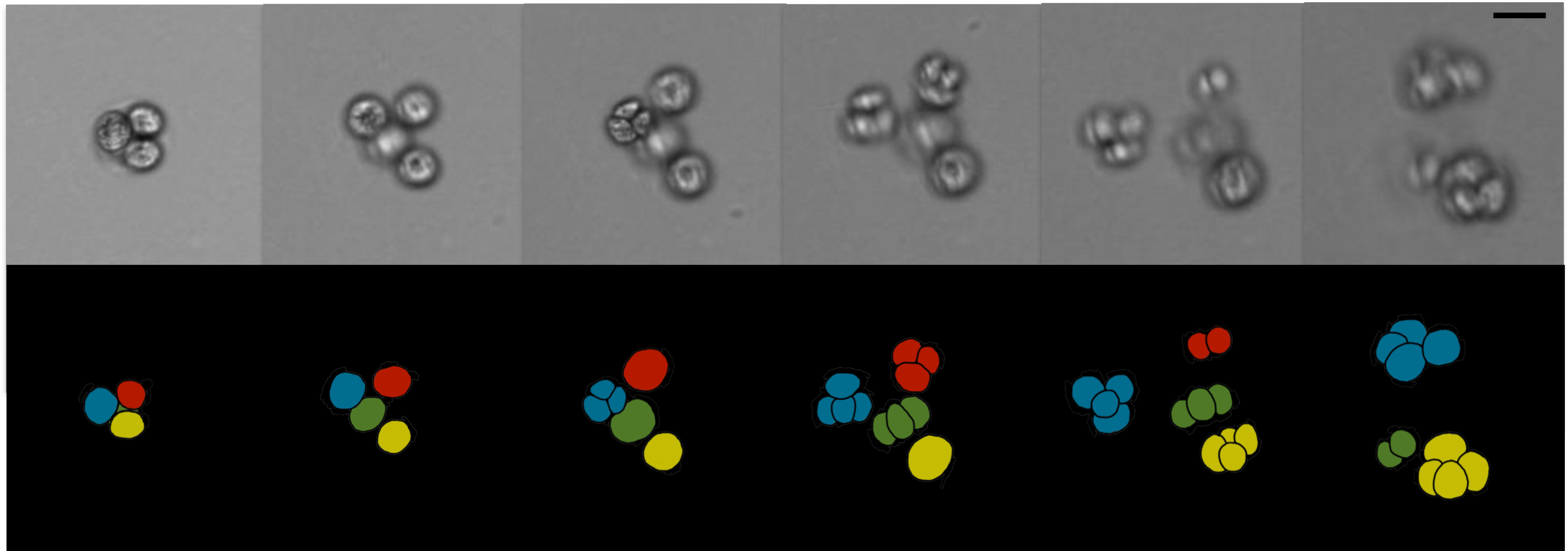
Evolved multicellular algae possess a multicellular lifecycle. Top row shows stop action stills of a video capturing the asexual reproduction of a single colony. Bottom row, graphical illustration of the same colony, with colors highlighting the daughter cell lineages derived from the cells within the mother colony. Scale bar equals 10µm.

## Discussion

The results of our experiments clearly match two theoretical expectations. The first [1] is that viscosity has a disproportional effect on motility in single cells compared with multicells due to the increase in collective thrust (58–60). As viscosity increases, we observe single cells decrease in speed while multicells do not (Fig. 3). The second [2] expectation met by our experiment is that as viscosity increases, the speed needed to maintain balance between nutrient acquisition and metabolism is equal to the speed of the ancestral unicell in the original low viscosity (55). Not only do we observe a disproportionate effect of viscosity on single cells and multicells, we observe that multicells maintain a speed in high viscosity which, as predicted (55), corresponds to the speed of single cells in 1x viscosity (Fig. 3).

The high viscosity of Snowball Earth oceans provided a universal selective barrier that could have been overcome using a variety of strategies. As both increased speed and decreased size are possible adaptations for high viscosity, it is not surprising to see these phenotypes — such as fast-moving ciliates (61) and unusually small alpha cyanobacteria (62) - arise at this same time in Earth history. Other eukaryotic adaptive strategies for motility and nutrient acquisition likely evolved during this time as well (45, 48). Many marine macroalgae rely on the surrounding current to bring nutrients to their blades (63); high seawater viscosity may explain the observed increase in surface area to volume ratio in the Cryogenian (20).

The idea of “complex multicellularity” tends to be entirely phenotypic — defined by large size, division of labor, and spatial organization (2, 64). Although none of the individual phenotypes we evolved in the lab possess all three aspects of complex multicellularity (2, 64), at least two are present independently. Large size is apparent in colonies visible to the naked eye, and spatial organization is present in motile multicellular forms with externally oriented flagella and coordinated behavior. The third aspect of complexity — division of labor – could potentially be present as well. The pre-existing variation of *C. reinhardtii’s* metabolism could be utilized evolutionarily, differentiating metabolically in response to nutrient availability or position within a large three-dimensional colony of cells (64–66). Moreover, the flagellar constraint on mitosis (64, 65, 67, 68) may also provide an opportunity for functional division of labor to evolve.

By this phenotypic definition, we observe “complexity” arise multiple times through Earth history (for example in *Bangiomorpha*, *Proterocladus*, and *Qingshania*). But there is a macroevolutionary facet of complex multicellularity exemplified by dominant modern clades that possess it—that of evolutionary success. Our experiment couples the phenotypic aspects of multicellularity with a relevant environmental pressure in which multicellularity not only evolves, but also provides an advantage to being complex. So why did modern multicellular lineages take so long to diversity if multicellularity is so easy to evolve? Our framework helps to resolve this paradox. The environment to globally select for complex life maybe just didn’t exist until the Snowball Earth.

Traits we observe to evolve in response to high viscosity, such as heritable multicellularity and organized flagellar orientation, would have also been advantageous in the warm, post-glacial oceans. As glaciers melted at the end of the Cryogenian period, nutrients were rapidly released into coastal waters (69). The resulting eutrophic environments would have led to a burst in primary productivity (39). In this competitive environment, the novelty of becoming larger and moving faster in order to survive the cold, viscous conditions of Snowball Earth oceans would lead to an ecological innovation in the post-Snowball world. Nascent motile multicellular lineages could take advantage of low marine viscosities to manipulate their surroundings (55), feeding in new and previously inaccessible ways. The ability to control the local environment — as opposed to being subject to it — opened the door for the origin and specialization of the dominant multicellular lineages on Earth today.

## Materials and Methods

### Model Organism

Before the experiment began, we grew wild-type *C. reinhardtii* CC-125 from the Chlamydomonas Resource Center in 10ml cultures of liquid HSA medium to an optical density (OD750) of 0.7 (∼2−10 6 cells/mL). We used a 500 nm light plate shaking at 100 rpms to encourage motility. Culture life cycles were not synchronized as we wanted diversity of morphologies present when inoculated into the experimental plate. Four celled colonies are observable during division, but due to the flagellar constraint on division and the presence of the mother cell wall, outward facing flagella that allow for motility are rare. The typical size of a single cell of this strain is around 10 µm in diameter (volume of about 270 µm3) and we observed speeds of motile individuals ranging from 10 to 40 µm s-1.

### Evolution Experiment

We designed a large petri plate (24.5 cm in diameter) with radial step changes of increasing viscosity (3.5 cm ring width). Ficoll400 was added to 1.5% agar HSA medium to create a solid base layer with different viscosities in each ring. The hard agar base layer was covered with 3 mm of HSA as a swim layer. We waited three days for the Ficoll400 to diffusively equilibrate between the hard agar layer below and the swim later above. The plate maintained stable sections of 1x, 2x, and 4x viscosities relative to water at room temperature (25 °C). We placed the plate on to a 500 nm light plate to equilibrate, and when it was ready, 100µL of culture was used as the inoculant in the center of the plate. *C. reinhardtii* grew for 30 days, corresponding to ∼70 generations (10.5-hour gen time) when the population reached the ring of media with 4x the relative viscosity of HSA media. We used a Multisizer 4s Coulter Counter to quantify size distributions and light microscopy along with custom motion tracking MATlab code to view and quantify motility.

### Phenotypic Plasticity Experiments

As a comparison, we cultured wild type *C. reinhardtii* that had been directly inoculated into the higher viscosity in order to quantify changes due to phenotypic plasticity.

Nine 15 mm petri dishes were set up on a 500 nm light plate. Three plates were used for each of the viscosities (1x, 2x, 4x relative viscosities). Cultures grew for approximately 7 generations before 1 ml samples from each were taken for Coulter Counter and microscopy analysis.

### Media

We used HSA media from the Chlamydomonas Resource Center and added Ficoll400 in corresponding amounts (0% for 1x, 5.5% for 2x, 9% for 4x) to the solid agar base layer. A single swim layer of HSA media was poured over top to a depth of 3 mm. For the plasticity experiments, Ficoll400 was added directly to HSA with the same weight to volume percentages as listed above. Dynamic viscosities of both the macroplate and individual petri dishes used for plasticity were calculated to be 0.95cP for 1x, 2.3cP for 2x, and 4.1cP for 4x relative viscosities.

### Transfers post Experimental Plate

A transect from the plate with 1x, 2x, and 4x viscosities were transferred into 15 mm petri plates as well as 10 mL culture tubes with corresponding viscosities. Samples were also taken and placed into plates of 1x viscosity in order to observe the stability of phenotypes over time. The stability of the size distributions acts as a measure of the bulk heritability of all phenotypes. These cultures were transferred for ∼70 generations in the 1x viscosity before taking Coulter Counter measurements to compare with the wild type *C. reinhardtii* grown in its native 1x viscosity.

### Coulter Counter

A sample was taken from each viscosity and we quantified size distributions with a Beckman Coulter Counter Multisizer 4s. 200 µm (20 mL accuvette) and 1000 µm (100 mL beaker) aperture tubes were used with 50 µl and 100 µl of sample respectively to include the full range of sizes from single cells to large colonies. Only the 200 µm aperture tube was used for heritability measurements in order to be able to focus on motile multicellular individuals. Standard operating methods were set to count and measure the individuals in a set volume of 1 ml of sample and electrolyte solution.

### Microscopy

We quantify the speed of motile unicells and multicells in each viscosity by randomly selecting sections on a slide and recording 10 second videos under a light microscope. After cleaning the videos of background noise, we used cell tracking code on Matlab to identify size and speed. We counted the number of cells in multicell phenotypes directly from these motility videos.

## Acknowledgements

The authors would like to thank Adam Younkin for help fabricating the macroplate, Anton P. Avramov, Jeffrey Cameron, Joe Dragavon, and Jian Wei Tay for help with microscopy, Sarah Hurley, Sarah Leventhal, Jinchung Li, Pat Kocielek, and Rebecca Safran for helpful discussion and feedback. AH would like to thank the NSF GRFP.

## References

1. N. J. Butterfield, Modes of pre-Ediacaran multicellularity. Precambrian Research 173, 201–211 (2009).

2. A. H. Knoll, The multiple origins of complex multicellularity. Annual Review of Earth and Planetary Sciences 39, 217–239 (2011).

3. L. Miao, Z. Yin, A. H. Knoll, Y. Qu, M. Zhu, 1.63-billion-year-old multicellular eukaryotes from the Chuanlinggou Formation in North China. Science Advances 10, eadk3208 (2024).

4. W. C. Ratcliff, R. F. Denison, M. Borrello, M. Travisano, Experimental evolution of multicellularity. Proceedings of the National Academy of Sciences 109, 1595–1600 (2012).

5. R. Fisher, T. Bell, S. West, Multicellular group formation in response to predators in the alga Chlorella vulgaris. Journal of evolutionary biology 29, 551–559 (2016).

6. S. E. Kapsetaki, R. M. Fisher, S. A. West, Predation and the formation of multicellular groups in algae. Evolutionary Ecology Research 17, 651–669 (2016).

7. M. D. Herron et al., De novo origins of multicellularity in response to predation. Scientific reports 9, 2328 (2019).

8. C. J. Rose, K. Hammerschmidt, P. B. Rainey, Experimental evolution of nascent multicellularity: Recognizing a Darwinian transition in individuality. BioRxiv (2020).

9. M. J. Hopkins, D. W. Bapst, C. Simpson, R. C. Warnock, The inseparability of sampling and time and its influence on attempts to unify the molecular and fossil records. Paleobiology 44, 561–574 (2018).

10. C. R. Marshall, The fossil record and estimating divergence times between lineages: Maximum divergene times and the importance of reliable phylogenies. Journal of Molecular Evolution 30, 400–408 (1990).

11. M. Foote, J. Hunter, C. Janis, J. J. Sepkoski, Jr, Evolutionary and preservational constraints on origins of biologic groups: divergence times of eutherian mammals. Science 283, 1310 (1999).

12. G. E. Budd, R. P. Mann, Two notorious nodes: a critical examination of MCMCTree relaxed molecular clock estimates of the bilaterian animals and placental mammals. bioRxiv, 2022.2007. 2001.498494 (2022).

13. M. Pennell (2023) Genes are often uninformative for dating species’ origins. (Nature Publishing Group UK London).

14. M. dos Reis et al., Uncertainty in the timing of origin of animals and the limits of precision in molecular timescales. Current Biology 25, 2939–2950 (2015).

15. D. B. Mills, W. R. Francis, D. E. Canfield, Animal origins and the Tonian Earth system. Emerging Topics in Life Sciences 10.1042/ETLS20170160, ETLS20170160 (2018).

16. G. Burzynski, T. A. Dececchi, G. M. Narbonne, R. W. Dalrymple, Cryogenian Aspidella from northwestern Canada. Precambrian Research 336, 105507 (2020).

17. N. A. Heim et al., Hierarchical complexity and the size limits of life. Proc. R. Soc. B 284, 20171039 (2017).

18. E. C. Yang et al., Divergence time estimates and the evolution of major lineages in the florideophyte red algae. Scientific reports 6, 21361 (2016).

19. A. Del Cortona et al., Neoproterozoic origin and multiple transitions to macroscopic growth in green seaweeds. Proceedings of the National Academy of Sciences 10.1073/pnas.1910060117, 201910060 (2020).

20. N. Bykova et al., Seaweeds through time: Morphological and ecological analysis of Proterozoic and early Paleozoic benthic macroalgae. Precambrian Research 350, 105875 (2020).

21. Q. Tang, K. Pang, X. Yuan, S. Xiao, A one-billion-year-old multicellular chlorophyte. Nature Ecology & Evolution 4, 543–549 (2020).

22. G. Li et al., Tonian carbonaceous compressions indicate that Horodyskia is one of the oldest multicellular and coenocytic macro-organisms. Communications Biology 6, 399 (2023).

23. J. Liu et al., Macroscopic fossils from the Chuanlinggou Formation of North China: evidence for an earlier origin of multicellular algae in the late Palaeoproterozoic. Palaeontology 66, e12685 (2023).

24. C. Niu et al., Chuaria and Tawuia fossils from∼ 1.0 Ga rocks in North China: Implications for a polyphyletic origin of Chuaria and a potential biological link between these two widespread Proterozoic taxa. Palaeogeography, Palaeoclimatology, Palaeoecology, 111966 (2023).

25. Q. Ye et al., The survival of benthic macroscopic phototrophs on a Neoproterozoic snowball Earth. Geology 43, 507–510 (2015).

26. J. Nursall, Oxygen as a prerequisite to the origin of the Metazoa. Nature 183, 1170 (1959).

27. K. M. Towe, Oxygen-collagen priority and the early metazoan fossil record. Proceedings of the National Academy of Sciences 65, 781–788 (1970).

28. E. A. Sperling et al., Oxygen, ecology, and the Cambrian radiation of animals. Proceedings of the National Academy of Sciences 110, 13446–13451 (2013).

29. A. H. Knoll, E. A. Sperling, Oxygen and animals in Earth history. Proceedings of the National Academy of Sciences 111, 3907–3908 (2014).

30. D. B. Mills et al., Oxygen requirements of the earliest animals. Proceedings of the National Academy of Sciences 111, 4168–4172 (2014).

31. D. B. Mills et al., The last common ancestor of animals lacked the HIF pathway and respired in low-oxygen environments. eLife 7, e31176 (2018).

32. E. A. Sperling, R. G. Stockey, The Temporal and Environmental Context of Early Animal Evolution: Considering All the Ingredients of an “Explosion”. Integrative and Comparative Biology 58, 605–622 (2018).

33. D. B. Cole et al., On the co-evolution of surface oxygen levels and animals. Geobiology (2020).

34. T. H. Boag, R. G. Stockey, L. E. Elder, P. M. Hull, E. A. Sperling, Oxygen, temperature and the deep-marine stenothermal cradle of Ediacaran evolution. Proceedings of the Royal Society B 285, 20181724 (2018).

35. R. G. Stockey, A. Pohl, A. Ridgwell, S. Finnegan, E. A. Sperling, Decreasing Phanerozoic extinction intensity as a consequence of Earth surface oxygenation and metazoan ecophysiology. Proceedings of the National Academy of Sciences 118, e2101900118 (2021).

36. E. A. Sperling et al., Breathless through Time: Oxygen and Animals across Earth’s History. The Biological Bulletin 243, 000–000 (2022).

37. C. Simpson, Coming together to understand multicellularity. Trends in Ecology & Evolution 38, 385–386 (2023).

38. G. O. Bozdag, E. Libby, R. Pineau, C. T. Reinhard, W. C. Ratcliff, Oxygen suppression of macroscopic multicellularity. Nature Communications 12, 2838 (2021).

39. J. J. Brocks et al., The rise of algae in Cryogenian oceans and the emergence of animals. Nature 548, 578 (2017).

40. C. T. Reinhard et al., Evolution of the global phosphorus cycle. Nature 541, 386–389 (2017).

41. M. Ingalls, J. Grotzinger, T. Present, B. Rasmussen, W. Fischer, Carbonate-Associated Phosphate (CAP) Indicates Elevated Phosphate Availability in Neoarchean Shallow Marine Environments. Geophysical Research Letters 49, e2022GL098100 (2022).

42. L. K. Eckford-Soper, K. H. Andersen, T. F. Hansen, D. E. Canfield, A case for an active eukaryotic marine biosphere during the Proterozoic era. Proceedings of the National Academy of Sciences 119, e2122042119 (2022).

43. K. J. Peterson, M. A. McPeek, D. Evans, Tempo and Mode of Early Animal Evolution: Inferences from Rocks, Hox, and Molecular Clocks. Paleobiology 31, 36–55 (2005).

44. S. M. Stanley, An ecological theory for the sudden origin of multicellular life in the late Precambrian. Proceedings of the National Academy of Sciences 70, 1486–1489 (1973).

45. S. Porter, The rise of predators. Geology 39, 607–608 (2011).

46. P. A. Cohen, L. A. Riedman, It’s a protist-eat-protist world: recalcitrance, predation, and evolution in the Tonian–Cryogenian ocean. *Emerging Topics in Life Sciences*, ETLS20170145 (2018).

47. D. B. Mills, The origin of phagocytosis in Earth history. Interface Focus 10, 20200019 (2020).

48. K. Dumack et al., It’s time to consider the Arcellinida shell as a weapon. European Journal of Protistology, 126051 (2024).

49. P. Branco, M. Egas, S. R. Hall, J. Huisman, Why do phytoplankton evolve large size in response to grazing? The American Naturalist 195, E20–E37 (2020).

50. M. LaBarbera, Precambrian geological history and the origin of the Metazoa. Nature 273, 22–25 (1978).

51. E. J. Trower, The enigma of Neoproterozoic giant ooids—Fingerprints of extreme climate? Geophysical Research Letters, e2019GL086146 (2020).

52. E. J. Trower, J. R. Gutoski, V. T. Wala, T. J. Mackey, C. Simpson, Tonian Low-Latitude Marine Ecosystems Were Cold Before Snowball Earth. Geophysical Research Letters 50, e2022GL101903 (2023).

53. Y. Ashkenazy et al., Dynamics of a Snowball Earth ocean. Nature 495, 90 (2013).

54. L. A. Riedman, P. M. Sadler, Global species richness record and biostratigraphic potential of early to middle Neoproterozoic eukaryote fossils. Precambrian Research 319, 6–18 (2018).

55. C. Simpson, Adaptation to a viscous Snowball Earth Ocean as a path to complex multicellularity. The American Naturalist 198, 590–609 (2021).

56. W. W. Crockett, J. Shaw, C. Simpson, C. P. Kempes, Physical constraints during Snowball Earth drive the evolution of multicellularity. bioRxiv, 2023.2012. 2007.570654 (2023).

57. M. Baym et al., Spatiotemporal microbial evolution on antibiotic landscapes. Science 353, 1147–1151 (2016).

58. B. Qin, A. Gopinath, J. Yang, J. P. Gollub, P. E. Arratia, Flagellar kinematics and swimming of algal cells in viscoelastic fluids. Scientific reports 5, 9190 (2015).

59. R. Podolsky, R. Emlet, Separating the effects of temperature and viscosity on swimming and water movement by sand dollar larvae (Dendraster excentricus). Journal of Experimental Biology 176, 207–222 (1993).

60. R. D. Podolsky, Temperature and water viscosity: physiological versus mechanical effects on suspension feeding. Science 265, 100–103 (1994).

61. N. M. Fernandes, C. G. Schrago, A multigene timescale and diversification dynamics of Ciliophora evolution. Molecular phylogenetics and evolution 139, 106521 (2019).

62. F. O. Aylward, J. C. Uyeda, C. A. Martinez-Gutierrez, A Timeline of Bacterial and Archaeal Diversification in the Ocean. bioRxiv (2022).

63. M. A. Koehl, Ecological biomechanics of marine macrophytes. Journal of Experimental Botany 73, 1104–1121 (2022).

64. J. T. Bonner, First Signals: The Evolution of Multicellular Development (Princeton University Press, Princeton, New Jeresey, 2001).

65. J. T. Bonner, On the origin of differentiation. Journal of biosciences 28, 523–528 (2003).

66. C. D. Schlichting, Origins of differentiation via phenotypic plasticity. Evolution & development 5, 98–105 (2003).

67. V. Koufopanou, The Evolution of Soma in the Volvocales. American Naturalist 143, 907–931 (1994).

68. R. E. Michod, D. Roze, Cooperation and conflict in the evolution of multicellularity. Heredity 86, 1–7 (2001).

69. P. F. Hoffman et al., Snowball Earth climate dynamics and Cryogenian geology-geobiology. Science Advances 3, e1600983 (2017).

70. M. Dohrmann, G. Wörheide, Dating early animal evolution using phylogenomic data. Scientific Reports 7, 3599 (2017).

71. N. J. Butterfield, Bangiomorpha pubescens n. gen., n. sp.: implications for the evolution of sex, multicellularity, and the Mesoproterozoic/Neoproterozoic radiation of eukaryotes. Paleobiology 26, 386–404 (2000).

72. J. Krissansen-Totton, R. Buick, D. C. Catling, A statistical analysis of the carbon isotope record from the Archean to Phanerozoic and implications for the rise of oxygen. American Journal of Science 315, 275–316 (2015).

73. T. M. Lenton, S. J. Daines, B. J. Mills, COPSE reloaded: An improved model of biogeochemical cycling over Phanerozoic time. Earth-Science Reviews 178, 1–28 (2018).

74. P. W. Crockford et al., Claypool continued: Extending the isotopic record of sedimentary sulfate. Chemical Geology 513, 200–225 (2019).

75. D. C. Catling, C. R. Glein, K. J. Zahnle, C. P. McKay, Why O2 Is Required by Complex Life on Habitable Planets and the Concept of Planetary “Oxygenation Time”. Astrobiology 5, 415–438 (2005).

76. G. Bell, The origin and early evolution of germ cells as illustrated by the Volvocales. The origin and evolution of sex 7, 221–256 (1985).

77. V. Koufopanou, The evolution of soma in the Volvocales. The American Naturalist 143, 907–931 (1994).

