## Supplemental information for "Experimental Snowball Earth Viscosity Drives the Evolution of Motile Multicellularity"

**This PDF file includes:**

Supporting text
Figure S1 to S5
Table S1
Legend for Movie S1and S2
Legends for Datasets S1 to S3

**Other supporting materials for this manuscript include the following:**

Movie S1 and S2
Datasets S1 to S3

**Calculating diffusion time for Ficoll.**

The diffusion of a molecule in a medium is described by Fick's second law of diffusion, which relates the rate of diffusion to the concentration gradient and the diffusion coefficient. The diffusion of Ficoll 400 from the agar layer into the swim layer can be calculated using the diffusion equation:

$$\varphi(x,t) = \varphi_0 erf(x/(2\sqrt{(Dt)}))$$

where $\varphi_0$ is the initial concentration at the source, erf is the error function, x is the distance from the source, D is the diffusion coefficient of Ficoll400, and t is the time.

To estimate the time required for Ficoll to fully diffuse from a 1 cm thick layer of 1.5% agar into a 3 mm liquid layer above, we can assume that the diffusion is complete when the concentration of Ficoll in the liquid layer is negligible (close to zero). In this case, we define a threshold concentration of 0.1% of $\varphi_0$ as the limit of detection. Using a diffusion coefficient of 1.8 x $10^{-7}$ $cm^2/s$ for Ficoll 400 in 1.5% agar at room temperature (25°C), we can calculate the time required to reach the threshold concentration at a distance of 1.5 mm (half the liquid layer thickness) from the source:

0.001 $\varphi_0$ = $\varphi_0$ erf(1.5 / (2 sqrt(D t)))

erf(1.5 / (2 sqrt(D t))) = 0.001

1.5 / (2 sqrt(D t)) = 3.0902

t = (1.5 / (2 sqrt(D)))^2 / (3.0902)^2

t = 56.9 hours = 2.4 days

Therefore, it would take approximately 2.4 days for Ficoll 400 to diffuse from a 1 cm thick layer of 1.5% agar into a 3 mm liquid layer above, assuming a diffusion coefficient of 1.8 x $10^{-7}$ $cm^2/s$ at room temperature. From the time the liquid layer was poured until inoculation, the plate was given three days to ensure full diffusion of Ficoll.

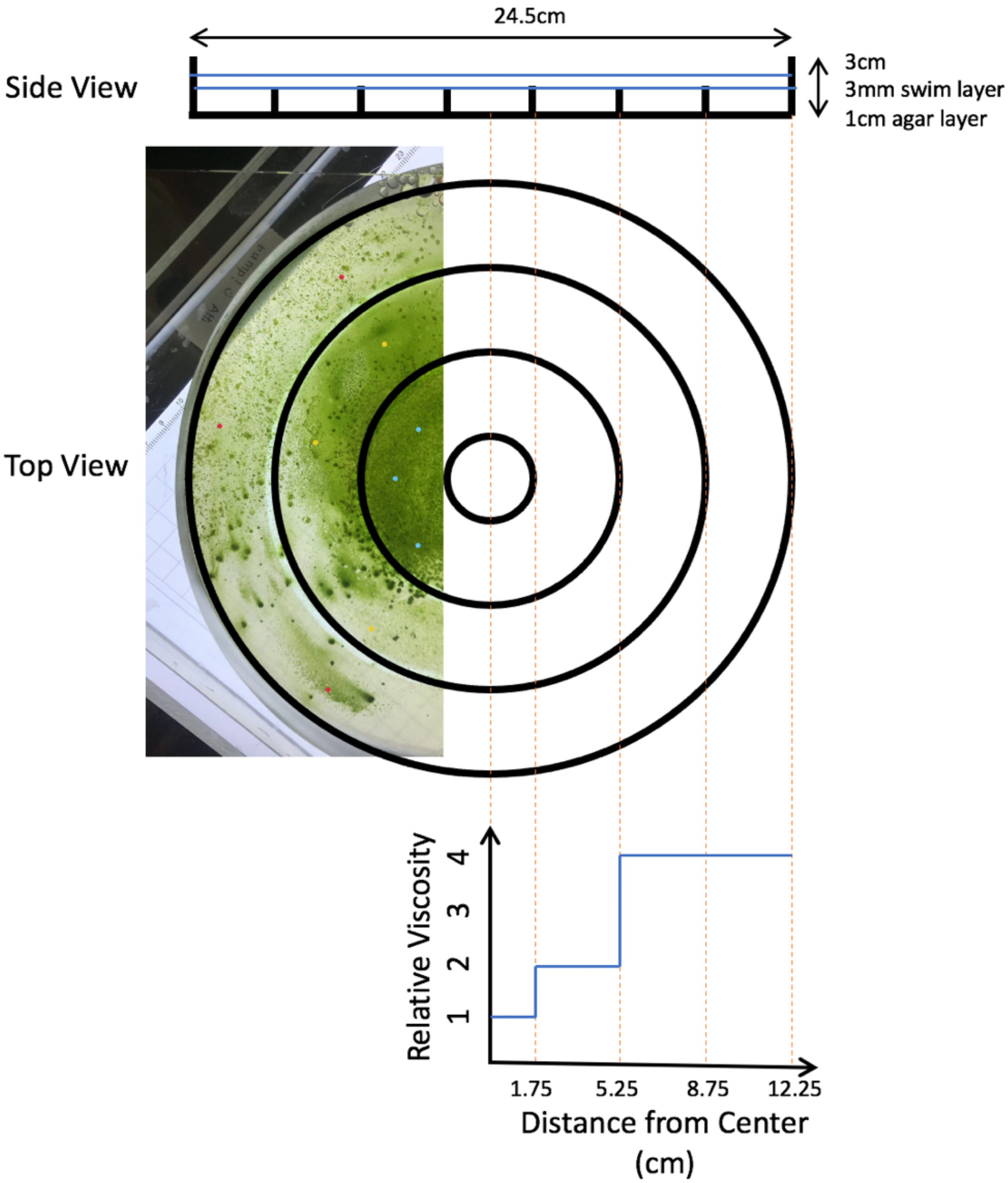


**Fig. S1.** Macro-plate experimental set-up from top and side views. The Macroplate, a 24.5 cm diameter petri dish, featured radial step changes in viscosity (3.5 cm ring width). Ficoll400 in 1.5% agar HSA medium created a solid base layer with varying viscosities. After three days of equilibration, the plate maintained stable sections with 1x, 2x, and 4x viscosities relative to water at 25 °C. *Chlamydomonas reinhardtii* culture (100 µL) was inoculated at the center, and growth over 30 days revealed distinct viscosity gradients, reaching 4x viscosity in the outer ring.

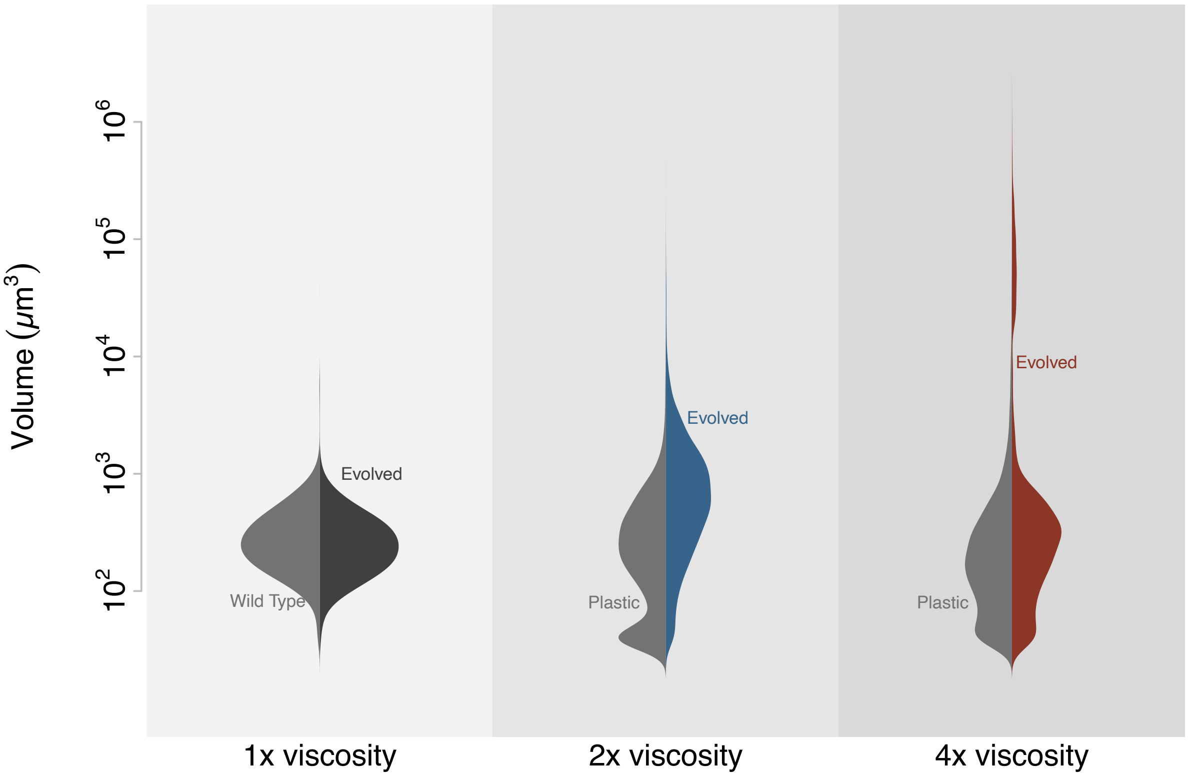


**Fig. S2.** Sample standardized distributions of evolved and plastic populations. Size distributions of *Chlamydomonas* cells were analyzed using a Coulter counter, comparing populations evolved in the Macroplate environment with those in a plastic control. Samples across aperture tubes have different sampling strategies, where large tubes measure a fixed volume of media where as small ones count specific numbers of organisms. Here we standardize sampling across aperture tubes to equal volumes of media. The violin plots illustrate the cell size frequencies for both populations, highlighting potential shifts or adaptations in cell size distribution due to the experimental conditions.

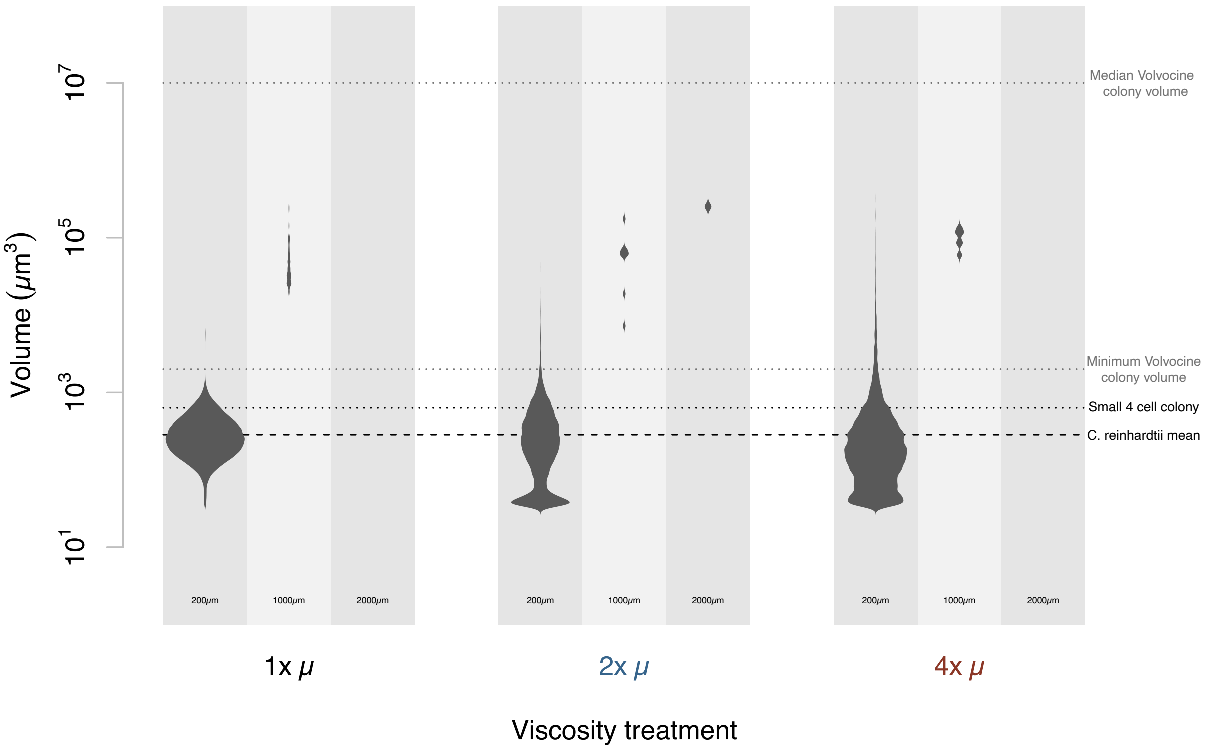


**Fig. S3.** Phenotypic plastic responses of *Chlamydomonas reinhardtii* populations directly inoculated in environments with varying viscosities. Size distributions, as quantified by Coulter Counter analysis, and microscopic observations reflect the observed phenotypes in response to media with 2x and 4x relative viscosities. Some large sizes are measured in the 1x viscosity as *C. reinhardtii* goes through multiple division during it's life cycle. Coordinated motility was not observed while typical stress-induced palmalloid phenotypes as well as small, non-motile single cells were present.

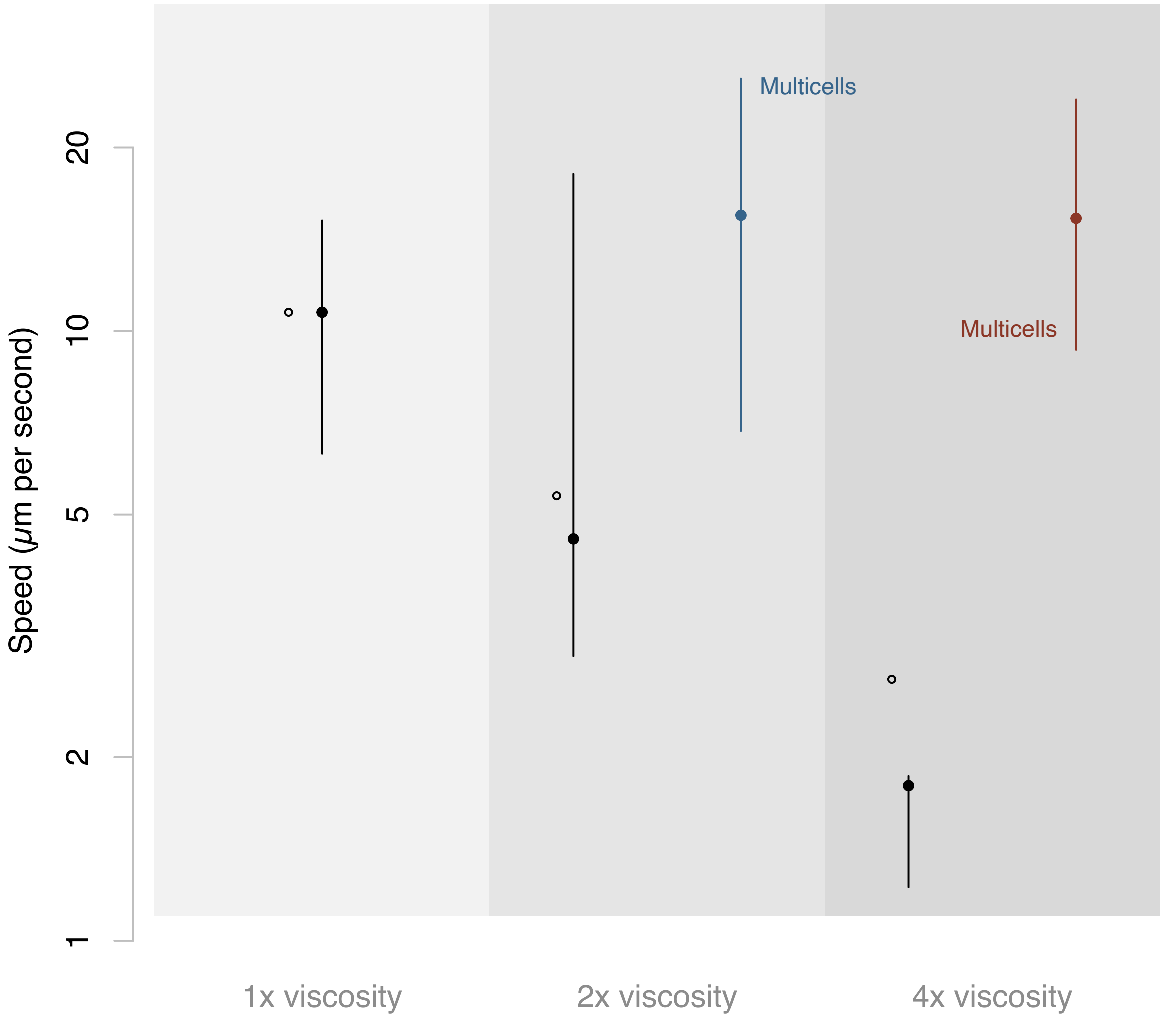


**Fig. S4.** Comparative analysis, including both observed speeds of evolved *Chlamydomonas reinhardtii* populations in environments with 1x, 2x, and 4x relative viscosities, as well as the predicted speeds of single cells in these viscous environments. The data showcase the expected decrease in speed for single cells as viscosity increases (based on Qin et al. 2015), aligning with theoretical predictions. The observed maintenance of higher speeds in multicellular colonies in 2x and 4x relative viscosities further supports the disproportionate impact of viscosity on unicells compared to multicells.

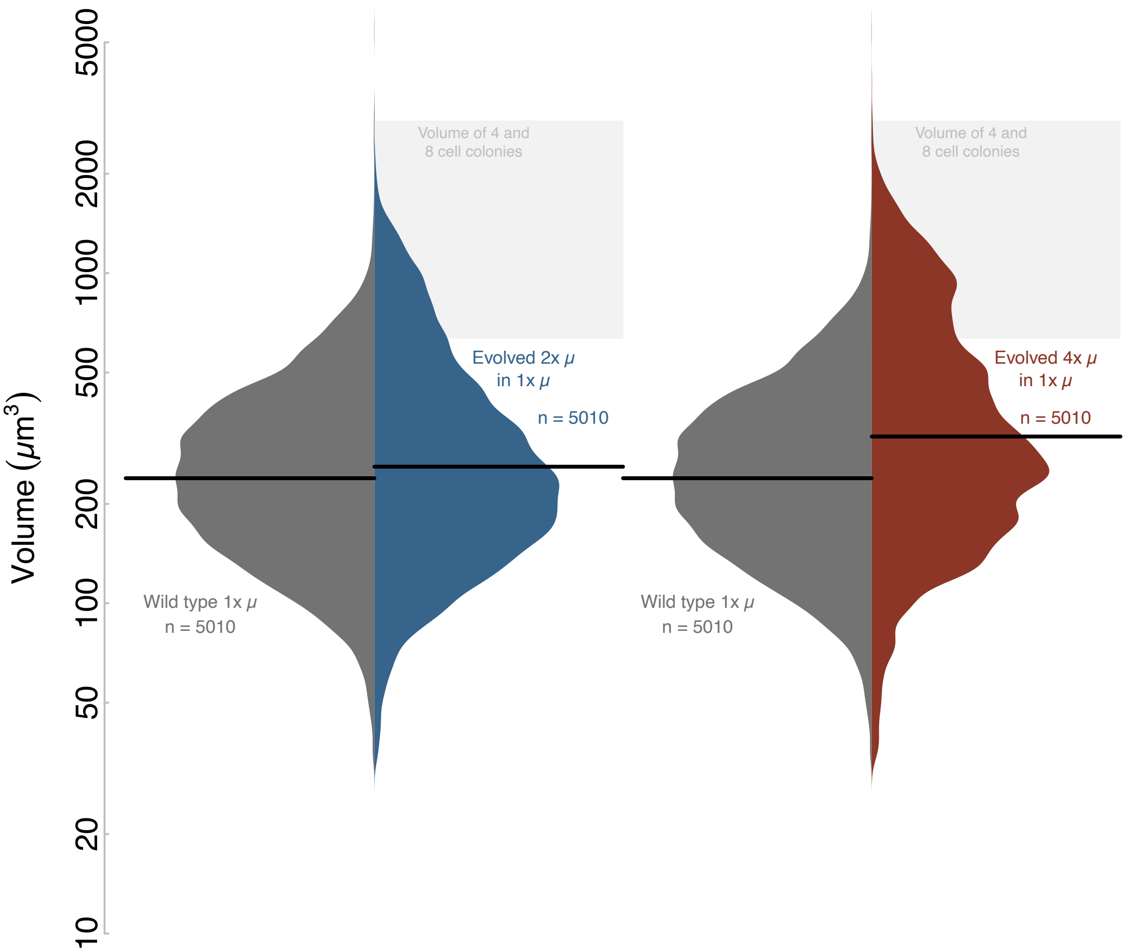


**Fig. S5.** Full distribution of heritability data of evolved *Chlamydomonas reinhardtii* populations. The data showcase the stability and heritability of size distributions across 1x, 2x, and 4x relative viscosities, providing insights into the bulk heritability of phenotypes over approximately 70 generations. The heritability measurements, obtained through Coulter Counter analysis, offer a quantitative assessment of the extent to which the evolved traits are passed on to successive generations. This analysis contributes to a comprehensive understanding of the persistence and inheritance of phenotypic characteristics under varying selective pressures imposed by viscosity gradients.

**Table S1.** Size ranges for each of the Coulter Counter Aperture Tubes used. Data obtained from beckman.com

| Aperture Tube | Diameter Range (µm) | | Volume Range (µm3) | |
|---|---|---|---|---|
| 200 | 4 | 160 | 33.5 | 905 x 10^3 |
| 1000 | 20 | 800 | 4189 | 113 x 10^6 |
| 2000 | 40 | 1600 | 33,510 | 905 x 10^6 |

**Movie S1 (separate file).** Evolved motile multicellular *C. reinhardtii*. Visual documentation of the evolved motile multicellular phenotypes of *Chlamydomonas reinhardtii* under light microscopy in response to increased viscosity. The film captures motility of a flagellated colony of eight cells, offering a visual representation of the evolved behaviors in the experimental setting.

**Movie S2 (separate file).** Time-lapse of evolved multicellular *C. reinhardtii* lifecycle. This footage shows various stages and behaviors exhibited throughout an asexual reproduction cycle, contributing to a nuanced understanding of life cycle dynamics under the selective pressures of viscosity.

**Dataset S1 (separate file).** SizeData.csv. Raw data of Coulter Counter size distributions from evolved and plastic populations.

**Dataset S2 (separate file).** SpeedData.csv. Raw data of speeds calculated from microscopy videos.

**Dataset S3 (separate file).** HeritabilityData.csv. Raw data of Coulter Counter size distributions from evolved populations placed back into low, 1x viscosity for 70 generations.
